# Genetic contributions to ectopic sperm cell migration in *Caenorhabditis* nematodes

**DOI:** 10.1101/302562

**Authors:** Janice J. Ting, Caressa N. Tsai, Asher D. Cutter

**Author notes:** corresponding author: Asher Cutter.

## Abstract

Reproductive barriers involving gametic incompatibilities can act to enhance population divergence and promote the persistence of species boundaries. Observing gametic interactions in internal fertilizing organisms, however, presents a considerable practical challenge to characterizing mechanisms of such gametic isolation. Here we exploit the transparency of *Caenorhabditis* nematodes to investigate gametic isolation mediated by sperm that can migrate to ectopic locations, with this sperm invasion capable of inducing female sterility and premature death. As a step toward identifying genetic factors and mechanisms associated with female susceptibility to sperm invasion, we characterized a panel of 25 *C. elegans* genetic mutants to test for effects on the incidence and severity of sperm invasion in both conspecific and interspecies matings. We found genetic perturbations to contribute to distinct patterns of susceptibility that identify ovulation dynamics and sperm guidance cues as modulators of ectopic sperm migration incidence and severity. Genotypes confer distinctive phenotypic sensitivities to the sperm from conspecific *C. elegans* males versus heterospecific *C. nigoni* males, implicating evolution of functional divergence in the history of these species for components of sperm-reproductive tract interactions. Sexually-antagonistic co-evolution within species that drives divergent trait and molecular evolution between species provides a working model to explain mismatched species-specific gametic interactions that promote or mitigate ectopic sperm migration.

**Article Summary:** Here we exploit the transparency of *C. elegans* roundworms to investigate reproductive barriers between species resulting from sperm-cell invasion into ectopic tissues, causing female sterility and premature death. We quantify female sensitivity to sperm invasion from conspecific and interspecific matings in a panel of 25 *C. elegans* genetic mutants, to demonstrate how ovulation dynamics and sperm guidance cues act as modulators of ectopic sperm-cell migration incidence and severity. We observe both conserved and divergent responses to different species, implicating evolution of functional divergence for components of sperm-reproductive tract interactions consistent with the outcome of sexually-antagonistic co-evolution.

## Introduction

Reproduction genetically binds individuals and populations to one another to define a species. Barriers to reproduction, therefore, will isolate populations from one another so that they accumulate genetic divergence as distinct lineages or species. Reproductive barriers that precede fertilization are especially potent because they inherently preclude offspring production, irrespective of whether zygotes would be viable or fertile (COYNE AND ORR 1989; COYNE AND ORR 1997). Courtship and mating behaviors allow pre-mating reproductive barriers to manifest at an early stage of the reproductive sequence (BOUGHMAN 2002; MAAN AND SEEHAUSEN 2011), yet in species with limited mate recognition, gametic interactions that occur during the post-mating pre-zygotic phase represent a critical period in the evolution of reproductive isolation (DOBZHANSKY 1951; EADY 2001; HOWARD *et al.* 2009). Given that gamete-related traits and the genes controlling them evolve rapidly (STOCKLEY 1997; SWANSON AND VACQUIER 2002; WILBURN AND SWANSON 2016), mismatched gamete interactions can evolve to create reproductive incompatibilities that impede the transfer of genetic material from one population to another to maintain, or foster formation of, distinct species (COYNE AND ORR 2004; HAERTY *et al.* 2007). Understanding the mechanisms and genetics that underpin the evolution of gametic reproductive isolation is therefore crucial to characterizing the speciation process (NOOR AND FEDER 2006; NOSIL AND SCHLUTER 2011).

Gametic reproductive incompatibilities between species are especially challenging to study with internal fertilization, though some recent advances provide novel views inside opaque organisms (MATTEI *et al.* 2015). Despite the difficulty in observing directly sperm and oocyte interactions inside the reproductive tract of a live female, studies show that sometimes heterospecific sperm are simply unable to outcompete conspecific sperm (i.e. ‘conspecific sperm precedence’), precluding formation of inter-species zygotes altogether (PRICE 1997; STOCKLEY 1997; EADY 2001). Even in the absence of sperm competition, however, transferred sperm or seminal products from heterospecific males can generate reproductive barriers between species by reducing female viability or fertility (PATTERSON 1946; KNOWLES AND MARKOW 2001; TING *et al.* 2014) or by disrupting intercellular interactions between sperm and egg (SNOOK *et al.* 2009).

The importance of such gametic barriers to overall reproductive isolation should be greater in organisms with weak pre-mating barriers like *Caenorhabditis* nematodes that often readily mate with other species (BAIRD 2001; GARCIA *et al.* 2007).

The transparent bodies of *Caenorhabditis* nematodes provide a convenient window for viewing gametic interactions (HILL AND L’HERNAULT 2001; HAN *et al.* 2010; MARCELLO *et al.* 2013; TING *et al.* 2014), providing a powerful testbed to screen for genetic factors that enhance or suppress gametic reproductive isolation between species. Normally, the amoeboid male sperm of *Caenorhabditis* crawl towards one of the paired spermathecae, where fertilization takes place, after insemination into the uterus via the vulva; the spermathecae represent the furthest points within the reproductive tract that male sperm ought to reach (Figure 1A) (MCCARTER *et al.* 1997; HUBBARD AND GREENSTEIN 2000). Interspecies matings between *Caenorhabditis* nematodes, however, often lead to a gametic form of reproductive isolation and reproductive interference: male sperm can cause sterility and reduced lifespan following matings between individual females or hermaphrodites to males from other species (TING *et al.* 2014). The heterospecific sperm not only displace any existing conspecific sperm from the sites of fertilization, but can migrate into ectopic meiotic and mitotic zones of the gonad, or even breach the reproductive tract altogether to meander in the body cavity (TING *et al.* 2014). This form of gametic isolation contrasts with the more widely-known pattern of conspecific sperm precedence in other animals (HOWARD *et al.* 2009). More rarely, sperm from conspecific males can migrate ectopically (TING *et al.* 2014). Although ectopic sperm invasion in *Caenorhabditis* exacts substantial harm to female physiology and fitness, distinct species pairs exhibit significant variation in both female susceptibility to sperm invasion and the relative degree of sperm mislocalization (TING *et al.* 2014). The genetic and mechanistic causes of this heterogeneity remain undetermined.

**Figure 1.**
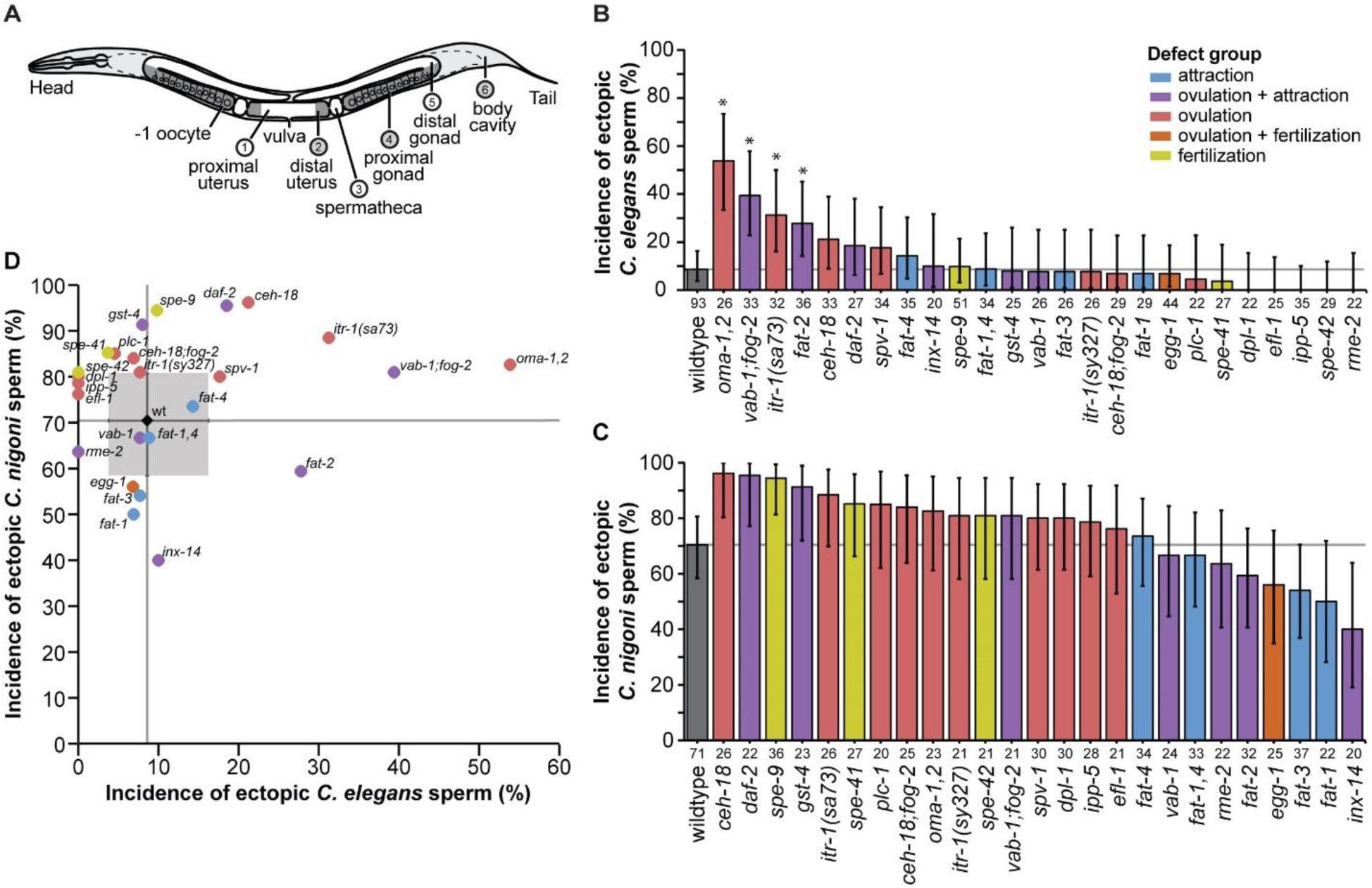
Genetic perturbations affect the incidence of sperm invasion in *C. elegans.* **(A)** Regions of *C. elegans* hermaphrodite body scored for sperm localization: Male sperm is transferred through the vulva into the uterus (regions 1-2), where the amoeboid sperm crawl to one of the paired spermathecae (region 3) which is the site of fertilization. Sperm are considered ectopic if found in the proximal gonad (region 4), distal gonad (region 5), and/or somatic locations outside the reproductive tract and gonad (region 6). **(B)** Incidence of ectopic sperm varies significantly across different *C. elegans* mutant strains (and wildtype strain N2) following mating with *C. elegans* males (% of hermaphrodites with ectopic sperm present in any location) (χ^2^=105.9, df=25, P<0.0001). **(C)** Incidence of ectopic sperm following heterospecific mating with *C. nigoni* males varies significantly among strains (χ^2^=76.2, df=25, P<0.0001). **(D)** The incidence of ectopic sperm across mutant genotypes from conspecific *C. elegans* males does not significantly predict the incidence of sperm invasion from heterospecific *C. nigoni* males (Spearman’s ρ= 0.21, df=24, p=0.3). Asterisks in (B) indicate statistical difference from wildtype after multiple test correction (Dunnett’s test α = 0.05). Sample sizes below bars in (B) and (C) indicate the number of mated *C. elegans* hermaphrodites scored; error bars show binomial 95% confidence intervals. Gray lines in (B-D) provide reference lines for wildtype values. Gray box in (D) indicates binomial 95% confidence intervals for the wildtype; error bars for mutants in (D) not shown for visual clarity. Mutant strains are colored to indicate phenotypic effects: ovulation defects (red), sperm attraction defects (blue), ovulation and attraction defects (purple), fertilization defects (yellow), ovulation and fertilization defects (orange), wildtype N2 (black) (Supplementary Table S1).

Here we test how genetic disruptions make the hermaphrodite (‘female’) sex of *C. elegans* more or less vulnerable to ectopic migration of male sperm cells of both their own species and of a different species (*C. nigoni*). *C. elegans* hermaphrodites experience high incidences of ectopic sperm after mating with *C. nigoni* males, but only rarely from males of their own species (TING *et al.* 2014). We predict that ectopic migration of heterospecific sperm arises as a direct consequence of the lack of sexual co-evolution between females and males of distinct species, whereas sexual selection within each species leads to female reproductive tracts with structural or signaling features compatible only with sperm from conspecific males. Specifically, co-evolved chemical signaling cues between gametes are hypothesized modulate susceptibility to interspecific ectopic sperm migration (TING *et al.* 2014). Mature oocytes in *Caenorhabditis* hermaphrodites and females secrete chemical cues that guide sperm to the sites of fertilization (reviewed in HAN *et al.* 2010; HOANG *et al.* 2013); we propose that the identity or sensitivity of signaling molecules, receptors, or transduction could differ among species, with ‘miscommunication’ resulting in ectopic migration of sperm cells. To address the mechanisms of ectopic sperm migration, we use a reverse genetics approach to identify genes important in the process of sperm invasion, analyzing the phenotypic effects of 25 genetic disruptions to understand the factors underlying ectopic sperm incidence and severity. We selected mutant strains to inform our hypotheses about potential roles of their deficient gene products in the fertilization process, based on previous studies demonstrating their influence on normal reproduction in terms of gamete signaling (HAN *et al.* 2010) and ovulation dynamics (KIM *et al.* 2013). By assaying sperm invasion incidence and severity in genotypes with perturbed sperm-oocyte signaling, spermathecal constriction/dilation responses, and oocyte maturation, we demonstrate how these factors influence the propensity for heterospecific sperm to migrate to ectopic locations within females and thus modulate the strength of reproductive isolation barriers between species.

## Methods

### Genetically disrupted *C. elegans* strains

To explore possible mechanisms associated with female susceptibility to ectopic migration of male sperm, we selected 25 *C. elegans* mutants from published literature and wormbase.org phenotype descriptions that we hypothesized might influence sperm invasion (Table 1; Supplementary Table S1). Phenotypic disruptions of the mutants affected the functions of physical structures or activity in the reproductive tract, such as changes in gonadal sheath cell contractions, ovulation, and cell-cell communication of gametes (Supplementary Table S1). We predicted sperm invasion to be more likely when sperm are able to reach the spermathecae, if ectopic sperm migration proceeds into the gonad through the proximal spermathecal valve. Consequently, we predicted that mutations that disrupt sperm guidance would reduce the likelihood of sperm invasion whereas mutations that disrupt proper sheath cell and spermathecal contractions may exacerbate ectopic sperm migration. In our assays, mutant strains otherwise share the same N2 genetic background, which acted as our wildtype control strain. Two strains required picking appropriate genotypes and phenotypes to assay from stocks that needed propagation as heterozygotes (TX183, SL1138). We maintained *Caenorhabditis* populations on 55 mm diameter NGM Lite agar plates with *Escherichia coli* (OP50) as food, using an agar concentration of 2.2% to discourage burrowing (STIERNAGLE 2006).

**Table 1.** Mutant *C. elegans* genotypes assayed for ectopic sperm presence.

| GENE (ALLELE)* | STRAIN | PROTEIN | PHENOTYPE GROUP** |
| --- | --- | --- | --- |
| wt (+) | N2 | n/a | wildtype |
| *ceh-18 (mg57)* | GR1034 | homeodomain transcription factor | ovulation |
| *ceh-18 (mg57); fog-2 (q71)* | DG1604 | homeodomain transcription factor | ovulation |
| *daf-2 (e1370)* | CB1370 | insulin/IGF receptor tyrosine kinase | attraction + ovulation |
| *dpl-1 (n3316)* | MT9940 | DP transcriptional regulator | ovulation |
| *efl-1 (n3639)* | MT11691 | E2F transcription factor | ovulation |
| *egg-1 (tm1071)* | AD186 | LDL-receptor | fertilization + ovulation |
| *fat-1 (wa9)* | BX24 | omega-3 fatty acyl desaturase | attraction |
| *fat-1 (wa9); fat-4 (wa14)* | BX52 | omega-3 and delta-5 desaturases | attraction |
| *fat-2 (wa17)* | BX26 | delta-12 fatty acyl desaturase | attraction + ovulation |
| *fat-3 (wa22)* | BX30 | delta-6 fatty acid desaturase | attraction |
| *fat-4 (wa14)* | BX17 | delta-5 fatty acid desaturase | attraction |
| *gst-4 (ok2358)* | RB1823 | GSH-dependent prostaglandin D synthase | attraction + ovulation |
| *inx-14 (ag17)* | AU98 | innexin gap junction channel | attraction + ovulation |
| *ipp-5 (sy605)* | PS3653 | inositol 5-phosphatase | ovulation |
| *itr-1 (sa73)* | JT73 | inositol (1,4,5) triphosphate receptor | ovulation |
| *itr-1 (sy327)* | PS2368 | inositol (1,4,5) triphosphate receptor | ovulation |
| *oma-1 (zu405te33); oma-2 (te51)* | TX183 | TIS11 zinc finger protein | ovulation |
| *plc-1 (rx1)* | PS4112 | phospholipase C | ovulation |
| *rme-2 (b1008)* | DH1390 | LDL-receptor | attraction + ovulation |
| *spe-41 (sy693)* | PS4330 | Ca<sup>+</sup> TRPC channel | fertilization |
| *spe-42 (tn1231)* | SL1138 | novel 7-pass transmembrane protein | fertilization |
| *spe-9 (hc88)* | BA671 | EGF repeat-containing protein | fertilization |
| *spv-1 (ok1498)* | RB1353 | F-BAR/RhoGAP protein | ovulation |
| *vab-1 (dx31)* | CZ337 | ephrin receptor | attraction + ovulation |
| *vab-1 (dx31); fog-2 (q71)* | DG1612 | ephrin receptor | attraction + ovulation |

\*genotype of full genetic background in Supplementary Table S1; \*\*ovulation=defects in sheath cell development or contractions, ovulation, oogenesis or oocyte maturation, sperm sensing, endomitotic oocytes, spermathecal valve constriction or contractions; attraction=defects in fatty acid synthesis and sperm migration toward oocytes; fertilization=defects in fertilization upon gamete contact

### Mating assays to quantify sperm invasion

We assessed the extent of ectopic sperm migration using three responses: whether any sperm occurred in ectopic locations (incidence), the number and identity of ectopic locations (severity), and how many sperm occurred in each location (abundance) (Figure 1A). To visualize and quantify the location and number of sperm transferred by males upon mating to *C. elegans* hermaphrodites (‘females’), we stained males of *C. nigoni* (strain JU1325) or *C. elegansfog*-*2(q71)* with MitoTracker^®^ Red CMXRos (Invitrogen) to fluorescently label their sperm (SCHEDL AND KIMBLE 1988; KUBAGAWA *et al.* 2006): batches of 300-600 males in 300 μL of M9 buffer with 10 μM MitoTracker^®^ Red CMXRos in a watch glass. We incubated males in the dark for two hours, then transferred males with a glass pipette to plates with food and allowed them to recover overnight before mating with age-matched virgin hermaphrodites. Following mating, the fluorescently stained sperm are visible inside the body of unstained hermaphrodites (TING *et al.* 2014).

To increase mating success, we immobilized *C. elegans* hermaphrodites in batches of 40-60 adult hermaphrodites anesthetized in a 0.2 mL tube with 80 μL of 0.1% tricaine and 0.01% tetramisole hydrochloride in M9 buffer for ~45 minutes (KIRBY *et al.* 1990; MCCARTER *et al.* 1997). Anesthesia does not affect sperm motility (KUBAGAWA *et al.* 2006). We then transferred hermaphrodites with a glass pipette onto a 55 mm diameter NGM-lite plate to allow the anesthetic to evaporate. Finally, we picked 10-20 individuals onto a 10 mm diameter bacteria spot (*E. coli* OP50) on a 35 mm diameter NGM-lite plate with stained males with a 1:3 hermaphrodite: male ratio. Males were allowed to mate for ~70 minutes, after which hermaphrodites were moved onto a fresh 35 mm diameter Petri dish with food and 20 μL of M9 buffer was added onto the hermaphrodites to aid in their recovery from the anesthesia (MCCARTER *et al.* 1999).

After mating, we quantified the location and abundance of sperm cells for each genotype that had been mated either to conspecific *C. elegans* males or to heterospecific *C. nigoni* males. First, we incubated mated hermaphrodites for 6 h at 20°C and then mounted them on 5% agarose pads on glass microscope slides, immobilized with 2 μL of 50 mM sodium azide (NaN_3_), and protected from rapid desiccation with a glass cover slip. Using an Olympus BX51 fluorescent compound microscope (40X magnification), we recorded the presence or absence of stained male sperm in six regions of the body (Figure 1A) (EDMONDS *et al.* 2011). Three regions represent non-ectopic zones (proximal uterus, distal uterus, and spermatheca), and three regions indicate ectopic sperm presence (proximal half of the gonad arm, distal half of the gonad arm, soma outside of the gonad and reproductive tract; Figure 1A). We quantified the ‘abundance’ of sperm localized to each region in each individual as: 0) ‘none’, no sperm in the region, 1) ‘low’, 1-5 sperm, 2) ‘medium’, <50 sperm (bright fluorescent signal, patchy distribution), and 3) ‘high’, ≥50 sperm (very bright fluorescent signal, continuous distribution). We consider the sperm invasion to be ‘severe’ whenever we observed any sperm outside of the gonad and reproductive tract (Figure 1), which was usually accompanied by sperm being present in other ectopic locations as well. Thus, the severity index for a given genotype corresponds to the percentage of mated hermaphrodites with ectopic sperm in locations outside the gonad and reproductive tract. We completed screening across 10 days in a one month period, where on each day N2 control matings were always present (n>4 for each conspecific and heterospecific matings). No block effect was detected for the wildtype strain (ANOVA: F_9, 154_=0.748, p=0.66), so we combined the data in subsequent analyses.

### Statistical analyses

We compared the incidence of sperm invasion for each *C. elegans* mutant genotype to the N2 wildtype when mated to either conspecific or heterospecific males, first by conducting an omnibus contingency table χ^2^-test for heterogeneity across the 26 strains and then applying a Dunnett’s test for binary data with multiple treatments and a single control (CHUANG-STEIN AND TONG 1995). In order to assess similarity of responses for the ‘amount’ of sperm localized to different regions of the body, we applied a clustering analysis separately for conspecific and heterospecific sperm, using the Bioinformatics Toolbox™ clustergram function in MATLAB™ (MATHWORKS™ 2017). Using sperm abundances (none, low, medium, high) in each region of the body (region 1-6; Figure 1), each genetic mutant generated a 24 dimensional data point (4 ‘amounts’ x 6 regions). We then calculated a Euclidian distance matrix between pairs of data points, which we treated as a similarity matrix that we then standardized by subtracting the mean and dividing by the standard deviation. Finally, we applied the UPGMA bottom-up hierarchical clustering method to create the agglomerative hierarchical dendrograms relating the similarity of sperm localization across genotypes.

### Data availability

Strains are available from the Caenorhabditis Genetics Center. The authors affirm that all data necessary for confirming the conclusions of the article are present within the article, figures, tables and supplementary files.

## Results

### Profiles of sperm cell invasion imply divergence in reproductive traits

To explore possible mechanisms associated with female susceptibility to ectopic migration of male sperm, we characterized 25 genetic disruptions in *C. elegans* that we hypothesized might influence sperm invasion. We screened the mutant strains of *C. elegans*, along with the wildtype N2 strain, for presence of ectopic male sperm in distinct regions of the body of mated hermaphrodites, contrasting animals that had mated to conspecific *C. elegans* males or to heterospecific *C. nigoni* males. We assessed the extent of sperm invasion in terms of overall ‘incidence’ as the fraction of animals with any ectopic sperm present, with its ‘severity’ measured by the number of ectopic locations with sperm present, which we also quantified in more detail with the ‘abundance’ of invasive sperm in each ectopic region.

Mutant strains differed significantly from one another in the incidence of ectopic sperm from both conspecific *C. elegans* males (χ^2^=105.9, df=25, P<0.0001) and from heterospecific *C. nigoni* males (χ^2^=76.2, df=25, P<0.0001). We found that wildtype *C. elegans* hermaphrodites had a substantially higher incidence of ectopic sperm when mated to heterospecific *C. nigoni* males (70% of individuals with ectopic sperm) than when mated to males of their own species (9%, χ^2^=67.315, df=1, P≤0.001; Figure 1; Supplementary Figure S1), consistent with previous work (TING *et al.* 2014). Mutant strains also exhibited more ectopic sperm overall in heterospecific matings than in conspecific matings (with the exception of *inx*-*14*; Figure 1; Supplementary Figure S1): heterospecific sperm were observed in ectopic locations in at least 40% of individuals across all genotypes, whereas just two mutant strains approached a comparable incidence of ectopic sperm migration from males of their own species (*oma*-*1*,*2* and *vab*-*1;fog*-*2;* Figure 1). Interestingly, the incidence of ectopic sperm did not correlate across genotypes for conspecific versus heterospecific matings (Spearman’s ρ=0.21, df=24, P=0.3; Figure 1), implying partial decoupling of how genetic perturbations confer sensitivity to conspecific versus heterospecific sperm.

Despite the relative rarity of ectopic sperm from conspecific males, four mutants showed a significantly higher incidence of sperm invasion compared to wildtype (*oma*-*1*,*2* 45% higher, *vab*-*1;fog*-*2* 31% higher, *itr*-*1(sa73)* 23% higher; *fat*-*2* 19% higher; Figure 1B). All four of these mutants exhibit ovulation-related defects, as would be predicted if the ability of sperm to migrate ectopically is influenced by structural or mechanical aspects of oocyte release. These observations suggest that the genetic pathways underlying the ovulatory process may be essential for female protection against the costs of mating with males of their own species.

When mated to heterospecific males, four other mutants showed ectopic sperm incidence that was >20% greater than wildtype (*gst*-*4*, *spe*-*9*, *daf*-*2*, and *ceh*-*18*) and two mutants gave values 20% less than wildtype *fat*-*1*, *inx*-*14*) (Figure 1C). Knockout of both *fat*-*1* and *inx*-*14* compromise sperm guidance to the spermathecae (KUBAGAWA *et al.* 2006; EDMONDS *et al.* 2011), so the tendency for ectopic migration to be reduced in these genetic backgrounds is consistent with sperm attraction to the spermathecae being crucial for the initiation of sperm invasion.

In some cases, we observed parallel effects of heterospecific and conspecific sperm on the severity of sperm invasion, such as for the three mutants that conferred the most extreme incidence of conspecific ectopic sperm that also tended to increase the incidence heterospecific sperm invasion (*itr*-*1(sa73)* 18% higher, *vab*-*1;fog*-*2* 11% higher, and *oma*-*1*,*2* 12% higher; Figure 1D). Similarly, the two mutants with most extreme incidence of ectopic heterospecific sperm also tended to increase the ectopic sperm incidence after conspecific matings (*daf*-*2* and *ceh*-*18*; Figure 1D). These parallel effects of both conspecific and heterospecific sperm imply that sperm invasion is controlled, in part, by overall sensitivity of the female reproductive tract to ectopic migration in similar ways to any source of sperm.

By contrast to these parallel effects of conspecific and heterospecific sperm, three mutant genotypes showed opposing trends of ectopic sperm migration. In the case of the *fat*-*2* mutant that disrupts oocyte secretion of sperm chemoattractants and exhibits spermathecal valve dilation (KUBAGAWA *et al.* 2006), we observed a pattern of higher conspecific but lower heterospecific sperm invasion than wildtype comparators (Figure 1D). Reciprocally, *spe*-*41* and *spe*-*42* showed a trend of reduced ectopic sperm incidence for conspecific sperm versus elevated incidence for heterospecific sperm (Figure 1D). These conflicting effects of conspecific versus heterospecific sperm imply that sperm invasion also is partly controlled by species-specific interactions of sperm with the reproductive tract, with contrasting outcomes for different pairings. Taken together, these findings support the idea that evolutionary divergence between species in some traits of male sperm and female reproductive tracts may modulate the propensity for ectopic sperm migration.

### Severity of sperm invasion is distinct from incidence

To assess in more detail the severity of how sperm invasion manifests, we quantified ectopic sperm presence in different regions of the hermaphrodite (‘female’) body. We scored severity based on the extent of spread of sperm through the body, ranging from contained within either the proximal or distal gonad, to spreading beyond the gonad into the body cavity (Figure 2A). In particular, we observed ‘severe’ ectopic migration by heterospecific *C. nigoni* sperm into the body cavity or somatic tissue outside the gonad altogether for nearly half of the 70% of wildtype *C. elegans* individuals that showed at least some degree of sperm invasion (i.e. severity = 46%; Figure 2). Across the 25 mutant genotypes, heterospecific ectopic sperm also often localized outside the gonad, leading to significant variation in severity scores across genotypes that ranged from 24% (*plc*-*1*) to 81% (*ceh*-*18;fog*-*2*) of individuals (χ^2^=67.97, df=25, P<0.0001; Figure 2C). Three mutants showed especially high severity, with heterospecific ectopic sperm occurring in the body cavity in 71% to 79% of mated individuals (*egg*-*1*, *spe*-*9*, *rme*-*2;* Figure 2C). By contrast, heterospecific sperm rarely invaded the body cavity for four *fat* mutants, showing unusually low severity of 27%-32% (Figure 2C). For those animals in which we found any ectopic heterospecific sperm, it most frequently localized to all three ectopic regions (proximal gonad, distal gonad, body cavity; Figure 2C).

**Figure 2.**
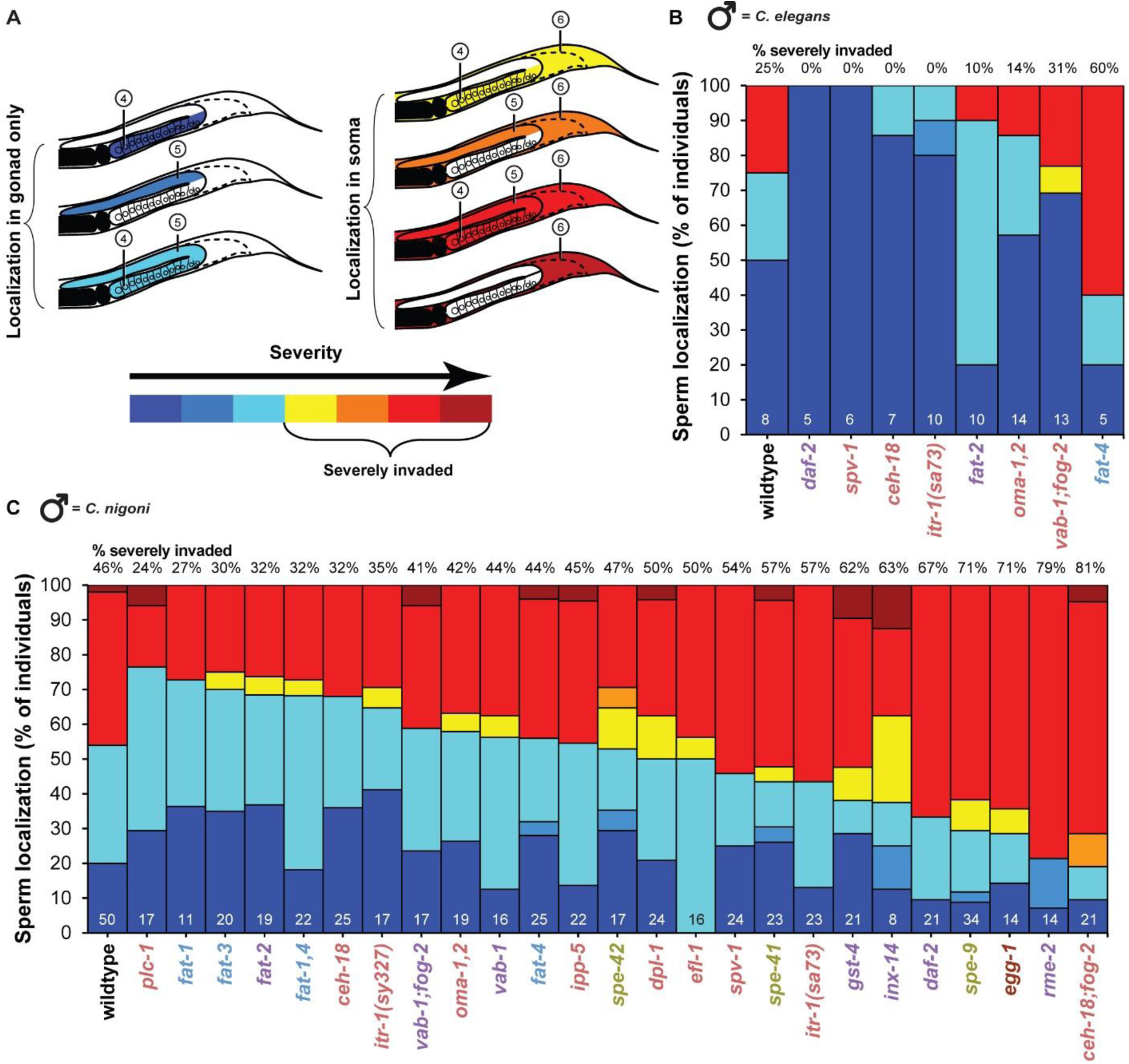
Sperm mislocalization in ectopic regions of *C. elegans* hermaphrodites. **(A)** Severity of sperm mislocalization was characterized for combinations of distinct ectopic regions, including the proximal and distal gonad arms as well as somatic zones of the hermaphrodite body outside of the reproductive tract altogether (see Figure 1). Individuals were considered ‘severely invaded’ when sperm was found in the soma and at least one other ectopic region. **(B)** Stacked bars indicate for eight different gene mutants the cumulative percentage of individual hermaphrodites with conspecific *C. elegans* male sperm mislocalized into the different ectopic regions. The other 17 mutant strains had fewer than five individuals with conspecific ectopic sperm, and were excluded from severity analysis. **(C)** Cumulative percentage of individual hermaphrodites with heterospecific *C. nigoni* male sperm mislocalized into the different ectopic regions of *C. elegans* hermaphrodites varies significantly across mutant genotypes (χ^2^=67.97, df=25, P<0.0001). Numbers above each bar in (B) and (C) indicate the percentage of severely invaded individuals among those with non-zero incidence of ectopic sperm. Numbers at the bottom of bars in (B) and (C) indicate sample size of individuals with ectopic sperm that allowed calculation of severity. Post-hoc statistical comparisons to wildtype identified no individual mutants after multiple test correction despite significant overall differences across strains (Dunnett’s test α = 0.05). Mutant strain names are colored to indicate functional phenotypic effects as in Figure 1. Sperm localization for non-ectopic regions is shown in Supplementary Figure S2.

By contrast to severity induced from heterospecific *C. nigoni* sperm, in those 9% of wildtype individuals that exhibited ectopic sperm from males of their own species, we found less severe localization patterns with the ectopic sperm cells localized solely to the proximal gonad 50% of the time, and with just 2 of the 8 individuals having sperm cells present outside the gonad (severity = 25%; Figure 2B). Indeed, mutant strains did not differ significantly from one another in degree of severe invasion of sperm from conspecific *C. elegans* males (χ^2^=33.8, df=25, P=0.11; Figure 2B).

Plots of ectopic sperm ‘incidence’ versus ‘severity’ highlight those mutant genotypes with unusual combinations of these two measures of sensitivity to sperm invasion relative to what is observed in wildtype individuals (Figure 3A). Incidence and severity provide complementary information about ectopic sperm migration, with their partial independence demonstrated with the lack of an overall correlation across genotypes between incidence and severity for both heterospecific and conspecific sperm invasion (Figure 3B). Nevertheless, we identified 10 mutants that had trends of both higher incidence of heterospecific sperm invasion and greater severity compared to wildtype, the most extreme examples being *ceh*-*18;fog*-*2*, *daf*-*2* and *spe*-*9* (Figure 3A). We also observed five mutants with the opposite trend, with tendencies for both lower severity and incidence of ectopic heterospecific sperm compared to wildtype, including four *fat* mutants (*fat*-*1*, *fat*-*2*, *fat*-*3*, and *fat*-*1*,*4:* Figure 3A).

**Figure 3.**
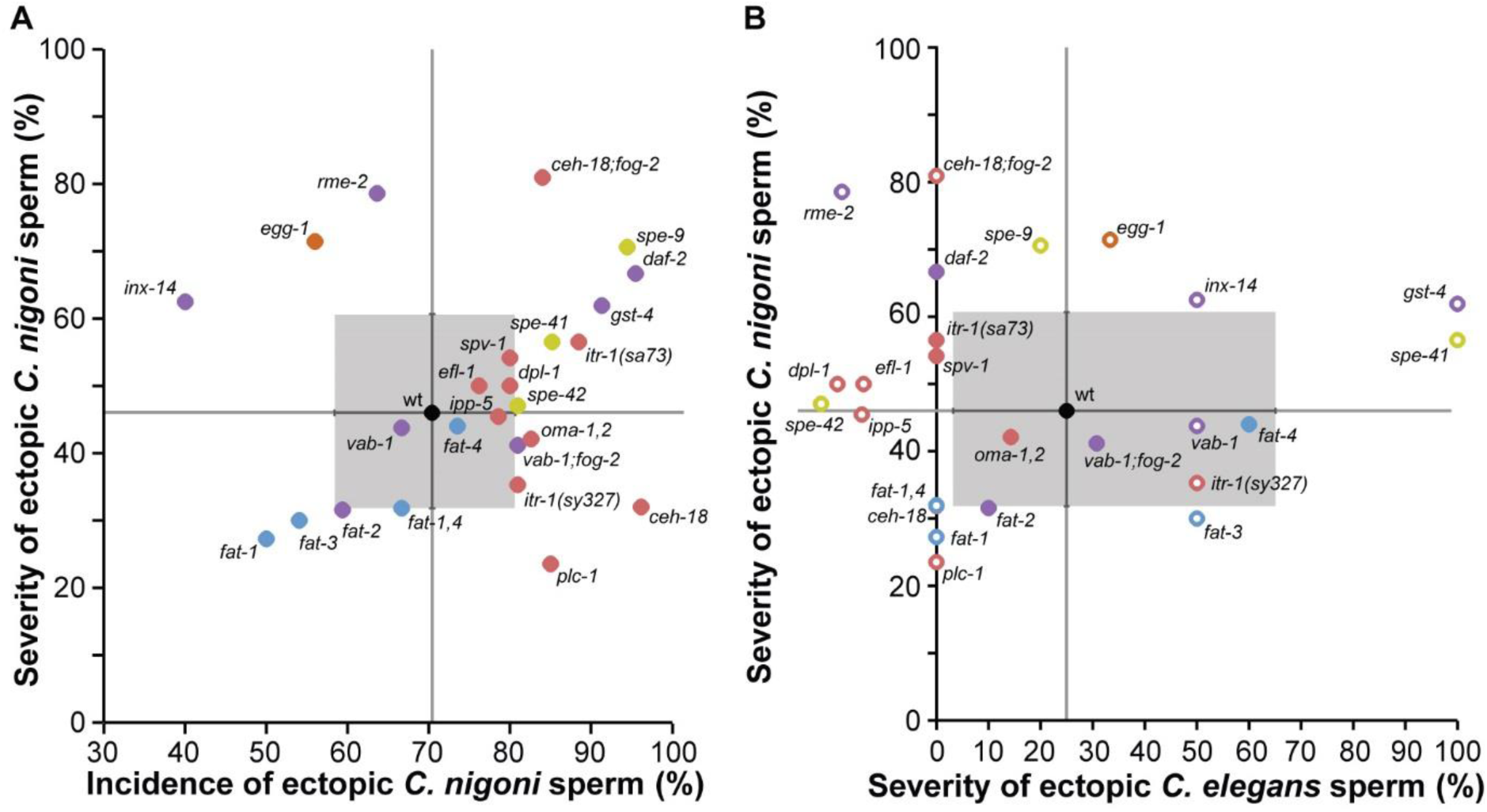
Severity of ectopic migration of sperm from *C. nigoni* and *C. elegans.* **(A)** The severity of ectopic sperm migration from *C. nigoni* males is partly decoupled from its incidence, showing no significant correlation across mutant genotypes (Spearman’s ρ= 0.21, df=24, P=0.31). Individual mutant strains with concordant influence on both severity and incidence occur in the upper-right and lower-left quadrants of severity × incidence space relative to the wildtype N2 strain values (wt). Mutant strain values in the upper-left and lower-right quadrants relative to wildtype exhibit discordant trends between incidence and severity when faced with heterospecific *C. nigoni* sperm. **(B)** The severity of ectopic sperm in *C. elegans* hermaphrodites exhibits species-specific outcomes for *C. nigoni* versus *C. elegans* males (Spearman’s ρ= 0.16, df=19, P=0.48). Gray lines provide reference lines for wildtype values; gray boxes indicate binomial 95% confidence intervals for the wildtype; error bars for mutants not shown for visual clarity. Error bars are larger for severity from *C. elegans* sperm due to the rarity of individuals that contained any ectopic sperm; mutant strains with <5 individuals observed to have ectopic sperm from *C. elegans* males are shown with unfilled points (mutants with zero individuals with ectopic sperm shown to the left of the y-axis). Mutant strains are colored to indicate phenotypic effects as in Figure 1 (Supplementary Table S1).

Interestingly, seven mutants exhibited a tendency for a higher incidence of sperm invasion and yet less severe locations of ectopic sperm occurrence, relative to wildtype (Figure 3A). Six of these seven genotypes show defects in spermathecal contractions and/or ovulation, the two most extreme examples being *plc*-*1* and *ceh*-*18* (Figure 3A). Notably, both of the *ceh*-*18* mutant genetic backgrounds that we analyzed exhibited unusual ectopic sperm invasion profiles, but in different ways: in a *fog*-*2* genetic background, *ceh*-*18* showed both high severity and incidence whereas the severity tended to actually be lower than wildtype when the *fog*-*2* gene was functional (Figure 3A). Lastly, we identified three mutants that tended to experience less sperm invasion than wildtype overall, yet exhibited more extreme severity in terms of ectopic locations (*rme*-*2*, *egg*-*1*, *inx*-*14*; Figure 3A). These disproportionate sensitivities to the incidence of ectopic sperm or to the severity of sperm invasion imply that these two aspects of ectopic sperm migration can be partly separated with distinct genetic perturbations to the female reproductive tract.

When we quantified sperm incidence in non-ectopic regions, we found both conspecific and heterospecific sperm most often to be present throughout both distal and proximal zones of the uterus and in the spermathecae (Figure 1A; Supplementary Figure S2).

### Genotype-phenotype clustering of sperm abundance profiles

Finally, we assessed the ‘abundance’ of sperm invasion by a semi-quantitative measure of the number of sperm (‘none’ to ‘high’ with ≥50 sperm cells) that localized to each of six regions of the body of mated individuals for each mutant genotype (Figure 4C). The distal uterus more often contained ‘high’ sperm abundance (both conspecific and heterospecific) compared to other regions, whereas no ectopic location exhibited ‘high’ sperm abundance in our assays 6 h post-mating, even for heterospecific sperm (Figure 4C; Supplementary Figure S3). We applied a clustering algorithm to group genotypes with similar profiles of sperm localization and abundance to distill these metrics into graphical summaries (Figure 4A-B).

**Figure 4.**
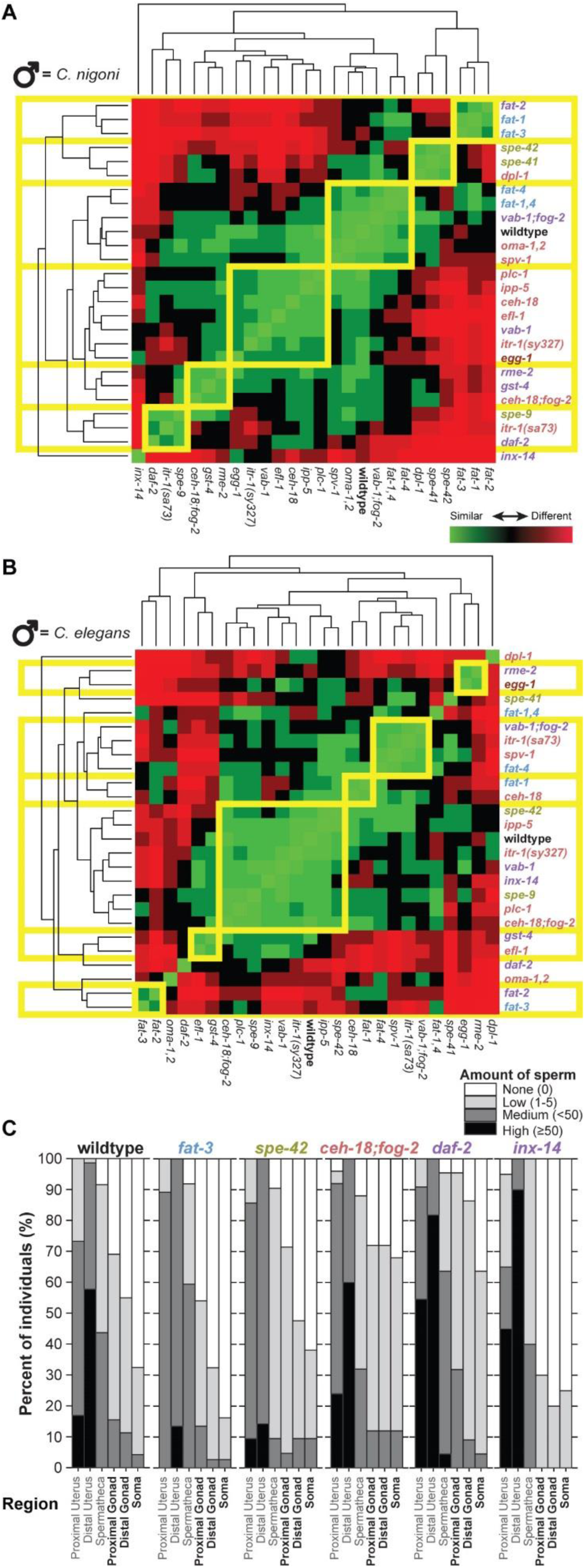
Phenotypic clustering of sperm abundance and distribution across mutant genotypes. **(A)** Hierarchical clustering groups together mutant strains with similar phenotypic responses in terms of the location and abundance of heterospecific *C. nigoni* male sperm in different body regions of hermaphrodite *C. elegans* (regions defined as in Figure 1). **(B)** Hierarchical clustering for profiles of sperm location and abundance for conspecific *C. elegans* male sperm in distinct body regions of hermaphrodites. Branchlengths in the dendrograms correspond to standardized phenotypic distance between mutant strains, which is depicted among all pairs of strains in the heat map as high similarity of sperm profile in green to high dissimilarity in red. Phenotypic clusters are outlined in yellow and mutant strain names are colored to indicate functional phenotypic effects as in Figure 1. **(C)** Stacked bars depict the cumulative relative abundance of heterospecific *C. nigoni* sperm observed in each region of the body as the proportion of mated hermaphrodites for wildtype strain N2 and five mutant strains (sperm abundances for all strains shown in Supplementary Figure S3; sperm abundances from conspecific matings for all strains shown in Supplementary Figure S4). Sperm abundance in each region for a given individual ranged from none (white) to high (≥50 sperm present, black), assessed for the three non-ectopic regions and the three ectopic regions defined in Figure 1. Mutant strain names are colored to indicate functional phenotypic effects as in Figure 1.

This clustering defined six distinct groups of genotypes when mated to *C. nigoni*, with one mutant strain exhibiting a unique profile characterized by especially low incidence and abundance of ectopic sperm (*inx*-*14*; Figure 4A). The five mutants that fell into the same cluster as the wildtype also occupied a similar region of interspecies severity × incidence space near the wildtype genotype (Figure 3A cf. Figure 4A). The three clusters of nine mutants (plus *inx*-*14*) that showed high dissimilarity to the wildtype also tended to be distant to wildtype in interspecies severity × incidence space, exhibiting especially high or especially low prevalence of ectopic sperm (Figure 3A cf. Figure 4A).

When mated to conspecific males, most mutants fell into two main clusters (Figure 4B). Eight mutants grouped with the wildtype profile of sperm localization and abundance, which included none of the *fat* mutants (Figure 4B). The nine mutants that were most dissimilar to wildtype did not fall into highly distinctive clusters of more than two strains for the conspecific sperm localization profiles (Figure 4B; Supplementary Figure S4). Overall, the multidimensional phenotypic clustering analysis reinforces the trends in ectopic sperm incidence and severity, while identifying affinities among mutant genotypes in sperm migration profiles not evident in those simple metrics (e.g. proximity of the heterospecific clusters containing *fat*-*3* and *spe*-*42*; Figure 4A).

## Discussion

We characterized the susceptibility of wildtype *C. elegans* and 25 mutants to sperm cell invasion: the incidence, severity, and abundance of sperm migration into ectopic locations beyond the reproductive tract. We captured the complementary contributions of these genes to distinct aspects of sperm migration by analyzing multiple metrics of sperm invasiveness, identifying factors related to species-specific sperm migration from the contrasting responses of mutant strains when mated to conspecific versus heterospecific partners. These experiments provide a crucial basis for understanding the mechanisms and genetics that underpin the evolution of gametic reproductive isolation (NOOR AND FEDER 2006; NOSIL AND SCHLUTER 2011), for *Caenorhabditis* nematodes in particular (TING *et al.* 2014). We found that genetic perturbations contribute to distinct responses of sperm invasion when sperm originated from either conspecific or heterospecific males. Genetic disruptions generally led to an elevated incidence of ectopic sperm localization, but some mutants that altered between-gamete communication instead tended to reduce the incidence and severity of ectopic sperm migration. We identify two mechanisms in particular that provide strong candidates for modulating the incidence and severity of ectopic sperm localization profiles: sperm chemical attraction defects lead to decreases, and ovulation defects increases, in the incidence and severity of sperm invasion.

Across genotypes, we observed that the incidence of ectopic sperm migration does not necessarily determine its severity. This result indicates that genetic perturbations to hermaphrodites (‘females’) can lead to independent consequences for i) how likely it is that male sperm will be able to migrate to ectopic locations at all and ii) how extensively different ectopic locations will be infiltrated. Moreover, the susceptibility of *C. elegans* hermaphrodites to sperm invasion from males of their own species or from another species (*C. nigoni*) also was uncorrelated across mutant genotypes overall. This finding of species-specific sensitivity to sperm invasion supports the idea that the distinct evolutionary trajectories of *C. elegans* and *C. nigoni* gave rise to divergent sperm × reproductive tract interactions, driving distinct hermaphrodite and female susceptibility to ectopic sperm migration. Theory predicts females to be more capable of resisting the costs of mating to males with which they co-evolved (PARKER AND PARTRIDGE 1998; PANHUIS *et al.* 2001; CHAPMAN *et al.* 2003). Sexually-antagonistic co-evolution within species that drives divergent trait and molecular evolution among species provides a prime possible explanation for the distinct phenotypic consequences quantified in our experiments (RICE 1996; MARKOW 1997; RICE AND HOLLAND 1997; STOCKLEY 1997; EADY 2001; ARNQVIST AND ROWE 2005).

### Sperm attraction defects and reduced sperm invasion

We were particularly interested in determining ectopic sperm profiles for mutants in the *fat* gene family, as these mutants are defective in conspecific sperm guidance by virtue of disrupting polyunsaturated fatty acid (PUFA) synthesis, the chemical precursors to oocyte-secreted F-class prostaglandin signaling molecules that direct sperm towards mature oocytes (WATTS AND BROWSE 2002; KUBAGAWA *et al.* 2006; HAN *et al.* 2010). If prostaglandin-based sperm chemotaxis is conserved across species, then we expected that the loss of these signals would lead to reduced invasion by both conspecific and heterospecific male sperm; previous research suggested the possibility that a high density of sperm in the spermatheca is a key precursor to ectopic migration (TING *et al.* 2014). Indeed, most *fat* mutants we tested tended to show both lower incidence and lower severity of sperm invasion by males of *C. nigoni* as well as *C. elegans* (*fat*-*1*,*fat*-*1;fat*-*4*, *fat*-*2*, *fat*-*3;* Figure 3A). These findings are consistent with sperm chemotaxis and sperm-oocyte signaling providing essential contributions to gametic isolation in *Caenorhabditis*.

We might further explore the individual ectopic sperm phenotypes of the *fat* mutants with an examination of the consequences of different mutations in the PUFA synthesis cascade. Briefly, *fat*-*1* mutants fail to produce omega-3 (n-3) PUFAs, *fat*-*2* mutants fail to produce Δ12-desaturase required to initiate PUFA synthesis, *fat*-*3* mutants lack Δ6 desaturase activity required to produce PUFAs, and *fat*-*4* mutants are defective in synthesizing Δ5 unsaturated fatty acids (WATTS AND BROWSE 2002; HAN *et al.* 2010). Moreover, disruption of *rme*-*2* leads to yolk accumulation in the pseudocoelom rather than getting transported to oocytes (GRANT AND HIRSH 1999), and yolk is where n-3 and n-6 PUFAs accumulate (KUBAGAWA *et al.* 2006). We hypothesize that this mislocalization of yolk that contains sperm chemoattractive cues leads to the pattern of extreme severity of heterospecific sperm invasion for *rme*-*2* mutants, but only when sperm have first migrated ectopically. Perhaps the endocytic trafficking of yolk from soma to germline represents a key vulnerability of females for ectopic sperm migration beyond the reproductive tract. We also found *fat*-*1* mutants to induce the greatest difference from wildtype when faced with sperm from heterospecific *C. nigoni* (Figure 3A), despite the lack of conspecific influence on sperm motility (KUBAGAWA *et al.* 2006); this result suggests that n-3 PUFA synthesis may be more crucial for *C. nigoni* sperm chemotaxis than for *C. elegans* sperm chemotaxis. Conversely, mutation to *fat*-*4* conferred the weakest effect on ectopic sperm migration, also eliciting minimal disruption to normal sperm taxis (KUBAGAWA *et al.* 2006), suggesting that Δ5 unsaturated fatty acids may be less important in regulating sperm chemoattraction and ectopic migration for both *C. elegans* and *C. nigoni.*

More generally, these findings suggest that divergence among species in the chemical constituents or stoichiometry of prostaglandin molecules and their PUFA precursors could play a key role in defining the likelihood and severity of ectopic sperm migration. Indeed, at least 10 chemically-related prostaglandin compounds collectively contribute to sperm guidance in *C. elegans* (HOANG *et al.* 2013), implicating ample scope for divergence across species. It remains an important goal to understand which components of sperm guidance may be conserved across species and which evolve species-specific roles. Recent work also shows that chemosensory cues from the external environment encountered by females and hermaphrodites, as well as their starvation state, can modulate female production of sperm chemoattractants that they secrete in their reproductive tract (KUBAGAWA *et al.* 2006; MCKNIGHT *et al.* 2014). Exogenous factors experienced by males also can influence sperm migration ability (HOANG AND MILLER 2017).

The evolution of unique combinations of sperm chemoattractants or environmental-dependence in different species could potentially contribute to the observed heterogeneity in ectopic sperm migration severity when distinct pairs of species interact (TING *et al.* 2014).

### Ovulation defects and elevated sperm invasion

In contrast to cell signaling-related defects, we predicted greater incidences of sperm invasion would result from ovulation defects. Ovulation normally begins with the contractions of gonadal sheath cells and the dilation of the distal spermathecal valve, which pulls the distal spermatheca over the most proximal mature (“-1”) oocyte (SCHEDL 1997). The distal part of the spermatheca subsequently constricts, holding the oocyte in the spermatheca for fertilization, followed by initiation of eggshell synthesis (WARD AND CARREL 1979; SCHEDL 1997). The fertilized egg then exits into the uterus within five minutes of the start of ovulation, aided by dilation of the valve between the spermatheca and the uterus (MCCARTER *et al.* 1999). In wildtype *C. elegans*, one oocyte is ovulated and fertilized at a time in an assembly line fashion, repeating every ~23 minutes (SCHEDL 1997; MCCARTER *et al.* 1999).

Our findings suggest that disruptions to different phases of the process of ovulation may enable ectopic sperm migration. For example, we observed severe invasion of heterospecific sperm in *itr*-*1(sa73)* mutants, which have been shown to exhibit continuous dilation and constriction of the spermatheca during ovulation, leading to oocyte tearing (BUI AND STERNBERG 2002; YIN *et al.* 2004). We propose that this process likely enhances the opportunity for sperm accumulated in the spermatheca to migrate into the proximal gonad if fragments of the torn oocyte prevent the spermatheca from completely constricting. Similarly, *fat*-*2* mutants are known to exhibit inappropriate spermathecal valve dilation during ovulation, resulting in misshapen eggs (KUBAGAWA *et al.* 2006; EDMONDS *et al.* 2010). In our experiments, *fat*-*2* mutants were more susceptible to conspecific but not heterospecific ectopic sperm than wildtype in spite of the sperm guidance cue defects also induced by *fat*-*2* (KUBAGAWA *et al.* 2006; EDMONDS *et al.* 2010), suggesting that maintenance of appropriate ovulation cues may be especially important for protection from ectopic conspecific sperm migration.

Regulation of spermathecal valve dilation and constriction is unlikely to be the only factor contributing to increased sperm invasiveness, however. For example, three mutants with defects in this process only exhibited unusual patterns of sperm invasion when mated to *C. nigoni* but not when mated to conspecific males (*daf*-*2*, *rme*-*2*, and *gst*-*4;* (GRANT AND HIRSH 1999; EDMONDS *et al.* 2010)). Moreover, gonadal sheath cell contractions distal to the spermatheca are slower than wildtype in *vab*-*1;fog*-*2* mutants, leading to delayed ovulation (MILLER *et al.* 2003), and ovulation from the gonad in *oma*-*1*,*2* mutants fails to occur altogether (DETWILER *et al.* 2001). Indeed, the incidence of ectopic sperm tends to positively coincide with the degree of ovulation defect among the four mutants we found to have individually significant increases in the incidence of conspecific sperm invasion (*fat*-*2*, *itr*-*1(sa73)*, *oma*-*1*,*2*, and *vab*-*1;fog*-*2*; Figure 1, Figure 3). The six mutants with the most extreme incidence and severity of sperm invasion from heterospecific *C. nigoni* males also commonly conferred ovulation defects (*itr*-*1(sa73)*, *daf*-*2*, *gst*-*4*, *ceh*-*18;fog*-*2*, *spe*-*41*, and *spe*-*9*; Figure 1, Figure 3).

Curiously, we observed contrasting patterns of sperm invasion in genetically distinct strains that both contained the *ceh*-*18(mg57)* mutation. *ceh*-*18(mg57)* confers delayed ovulation and slow, weak and uncoordinated sheath cell contractions compared to wildtype (ROSE *et al.* 1997), but one mutant strain also contained *fog*-*2(q71)* in the genetic background which eliminates self-sperm production (SCHEDL AND KIMBLE 1988). We observed nearly 50% more extreme sperm invasion for *ceh*-*18;fog*-*2* than for *ceh*-*18* alone (Figure 3). Because the *fog*-*2* genetic background prevents hermaphrodites from making their own self-sperm, this *fog*-*2*-dependent effect of sperm invasion on *ceh*-*18* mutants suggests that production of self-sperm may help to protect the gonad from ectopic migration of heterospecific sperm. We propose that such protection is a byproduct of self-sperm triggering ovulation and mechanical oocyte cell movement within the female germline prior to arrival of aggressive heterospecific sperm. Mutations to hermaphrodite sperm-associated genes *spe*-*41* and *spe*-*42* also lead to self-sterility, like *fog*-*2*, which may explain the tendency for elevated incidence and/or severity of ectopic sperm migration from both conspecific and heterospecific males for these mutants as well, although *spe*-*41* hermaphrodites ovulate similar to wildtype (XU AND STERNBERG 2003). Priming by self-sperm may only affect particular contexts, however, as we did not see much more ectopic sperm migration for either *vab*-*1* or *vab*-*1;fog*-*2* relative to wildtype (Figure 3), where VAB-1 is the ephrin receptor for the sperm-derived MSP trigger of ovulation (MILLER *et al.* 2003). We also observed unusually high incidence and severity of ectopic sperm migration in *spe*-*9(hc88)* mutants, which exhibit partial sperm-associated self-sterility at the semi-permissive temperature of 20°C used in our study (L’HERNAULT *et al.* 1988; SINGSON *et al.* 1998). The non-functional self-sperm nevertheless induce oocyte maturation and ovulation (SINGSON *et al.* 1998), however, so it remains unclear what mechanism might facilitate increased sperm invasion in *spe*-*9(hc88).* Disruption of *ceh*-*18* also leads to oocyte endomitosis (MCCARTER *et al.* 1997), such that oocytes fail to ovulate and undergo multiple rounds of endomitotic DNA replication, resulting in polyploid oocytes that remain in the gonad arm (IWASAKI *et al.* 1996). The accumulation of endomitotic oocytes might also lead to accumulation of secreted sperm-guidance cues and promote ectopic sperm migration.

Our screen of perturbed genotypes shows that disrupted ovulation can contribute to the incidence and severity of ectopic sperm migration from inter-species matings, but what implications does this have for natural variation and divergence among species? Species of *Caenorhabditis* differ substantially in ovulation rate, potential for self-sperm induced ovulation, egg retention and egg size (NIGON AND DOUGHERTY 1949; FARHADIFAR *et al.* 2015). Consequently, evolutionary change to ovulation dynamics across the phylogeny contributes a viable source of species differences in female susceptibility to ectopic sperm migration (TING *et al.* 2014).

### Male versus female contributions to divergent gametic interactions

Successful fertilization within *Caenorhabditis* is attributed largely to sperm chemotaxis, which we expect to be influenced by both oocyte signaling as well as sperm competitive ability. Our experiments focused on female factors that modulate sperm invasion, but genetic differences among males also are important to understanding ectopic sperm migration. Alleles of *comp*-*1*, *srb*-*13*, and *mss* genes that influence sperm competitive ability or guidance in *Caenorhabditis* provide clear candidates for exploring the sperm-oriented perspective (EDMONDS *et al.* 2010; HANSEN *et al.* 2015; YIN *et al.* 2018). Similarly, members of the *msp* multi-gene family are important in sperm signaling to oocytes (MILLER *et al.* 2001). For example, high concentrations of MSP induce sheath cell hypercontraction in *C. elegans* and the conserved 20 amino acids at the C-terminus of MSP exert cross-species capability of inducing sheath cell contractions (MILLER *et al.* 2001), suggesting that exceptional concentrations of MSP released into females by males of another species could promote ectopic sperm migration. Given the rapid evolution sperm traits and sequences of sperm-related genes (CUTTER AND WARD 2005; ARTIERI *et al.* 2008; VIELLE *et al.* 2016; YIN *et al.* 2018), we anticipate that such changes will prove important to understanding the variation among species in ectopic sperm migration patterns.

Unknown molecules on the sperm surface or in the seminal fluid might also enable sperm to breach membranes of the female reproductive tract. For example, sperm lysin protein in abalone creates a hole for sperm passage in the vitelline envelope surrounding the egg to allow fertilization (KRESGE *et al.* 2001). In *Ascaris* nematodes, the vitelline layer is present on oocytes in the oviduct (FOOR 1967), so penetration of a vitelline-like layer on the surface of the oocyte by *Caenorhabditis* sperm could predispose them to perforating other cell types. Sperm-egg fusion in *Caenorhabditis*, however, remains enigmatic (STEIN AND GOLDEN 2015), making it difficult to confidently ascribe the inherent properties of sperm cells in fertilization to their potential to penetrate other cell types. Parallels with pathogenesis also might be informative: the *Caenorhabditis* intracellular microsporidian parasite *Nematocida parisii* invades intestinal cells by traversing cell membranes (TROEMEL *et al.* 2008). Sperm cells from males of another species effectively act as a sexually-transmitted pathogen inside a foreign dead-end host (LEVIN AND BULL 1994; WOOLHOUSE *et al.* 2001). Future studies of what genetic factors promote or prevent sperm invasion from the perspective of sperm will be crucial for elucidating a full understanding of the causes and consequences of ectopic sperm migration in the evolution of gametic reproductive interference and reproductive isolation.

## Acknowledgements

We thank Jonathan Schneider for conducting the cluster analysis. We thank H. Robert Horvitz, from Massachusetts Institute of Technology, for providing strains and the CGC for providing strains (CGC funded by NIH Office of Research Infrastructure Programs: P40 OD010440). ADC is supported by funds from the Natural Sciences and Engineering Research Council of Canada.

## Author Contributions

Conceived and designed the experiments: JJT ADC. Performed the experiments: JJT CNT. Analyzed the data: JJT CNT ADC. Contributed reagents/materials/analysis tools: ADC. Wrote and edited the paper: JJT CNT ADC.

## Supplementary Tables

**Supplementary Table S1.** *C. elegans* mutant strains assayed for sensitivity to ectopic sperm migration after mating with males of *C. elegans* or *C. nigoni.*

| GENE | STRAIN GENOTYPE | STRAIN | PHENOTYPE GROUP | PROTEIN | LOCALIZATION | PHENOTYPE NOTES |
| --- | --- | --- | --- | --- | --- | --- |
| wt | wildtype | N2 | wildtype | n/a | n/a | n/a |
| *ceh-18* | *ceh-18(mg57) X* | GR1034 | **ovulation** | homeodomain transcription factor | broadly expressed | required for normal sheath cell development and contractions, ovulation, sperm sensing; loss induces endomitosis |
| *ceh-18; fog-2* | *ceh-18(mg57) X; fog-2(q71) V* | DG1604 | **ovulation** | homeodomain transcription factor | broadly expressed | required for normal sheath cell development and contractions, ovulation, sperm sensing; loss induces endomitosis |
| *daf-2* | *daf-2(e1370) III* | CB1370 | **attraction + ovulation** | insulin/IGF receptor tyrosine kinase | broadly expressed | required for normal sperm migration toward oocytes, spermathecal valve constriction |
| *dpl-1* | *dpl-1(n3316) unc-4(e120)/mnC1 [dpy-10(e128) unc-52(e444) II]* | MT9940 | **ovulation** | DP transcriptional regulator | broadly expressed | required for normal oogenesis, ovulation; loss induces endomitosis |
| *efl-1* | *efl-1(n3639) V/+ ; lin-15A(n767) X* | MT11691 | **ovulation** | E2F transcription factor | gonad | required for normal oogenesis, ovulation; loss induces endomitosis |
| *egg-1* | *egg-1(tm1071) III* | AD186 | **fertilization + ovulation** | LDL-receptor | oocyte membrane | required for normal gamete interaction and fertilization, ovulation |
| *fat-1* | *fat-1(wa9) IV* | BX24 | **attraction** | omega-3 fatty acyl desaturase | broadly expressed | PUFA synthesis |
| *fat-1,4* | *fat-4(wa14) fat-1(wa9) IV* | BX52 | **attraction** | omega-3 and delta-5 desaturases | broadly expressed |  |
| *fat-2* | *fat-2(wa17) IV* | BX26 | **attraction + ovulation** | delta-12 fatty acyl desaturase | broadly expressed | C18/C20 PUFA synthesis; required for normal sperm migration toward oocytes, spermathecal valve constriction |
| *fat-3* | *fat-3(wa22) IV* | BX30 | **attraction** | delta-6 fatty acid desaturase | broadly expressed | delta-6/C20 fatty acid synthesis |
| *fat-4* | *fat-4(wa14) IV* | BX17 | **attraction** | delta-5 fatty acid desaturase | broadly expressed | PUFA precursor synthesis |
| *gst-4* | *gst-4&msp-38(ok2358) IV* | RB1823 | **attraction + ovulation** | GSH-dependent prostaglandin D synthase | broadly expressed | required for normal sperm migration toward oocytes |
| *inx-14* | *inx-14(ag17) I* | AU98 | **attraction + ovulation** | innexin gap junction channel | broadly expressed | required for normal sperm migration toward oocytes, oocyte maturation |
| *ipp-5* | *ipp-5(sy605) X* | PS3653 | **ovulation** | inositol 5-phosphatase | distal spermatheca, proximal sheath cells | required for normal spermathecal contractions |
| *itr-1(sa73)* | *itr-1(sa73) IV* | JT73 | **ovulation** | inositol (1,4,5) triphosphate receptor | broadly expressed | required for normal ovulation, spermatheca and sheath cell contractions |
| *itr-1(sy327)* | *itr-1(sy327) unc-24(e138) IV* | PS2368 | **ovulation** | inositol (1,4,5) triphosphate receptor | broadly expressed | required for normal ovulation, spermatheca and sheath cell contractions |
| *oma-1,2* | *oma-1(zu405te33)/nT1 [unc-?(n754) let-?] IV; oma-2(te51)/nT1 V* | TX183 | **ovulation** | TIS11 zinc finger protein | broadly expressed | required for normal oocyte maturation, ovulation; no obvious single-mutant phenotypes |
| *plc-1* | *plc-1(rx1) X; kfEx2* | PS4112 | **ovulation** | phospholipase C | broadly expressed | required for normal ovulation, spermathecal contractions; loss induces endomitosis |
| *rme-2* | *rme-2(b1008)IV* | DH1390 | **attraction + ovulation** | LDL-receptor | near oocyte membrane | required for normal yolk uptake in oogenesis, sperm movement, ovulation, spermathecal valve constriction; loss induces endomitosis |
| *spe-41* | *spe-41(sy693) III; him-5(e1490) V* | PS4330 | **fertilization** | Ca<sup>+</sup> TRPC channel | sperm membrane | required for fertilization upon gamete contact |
| *spe-42* | *spe-42(tn1231) V/nT1 [unc-?(n754) let-? qIs50] (IV;V)* | SL1138 | **fertilization** | novel 7-pass transmembrane protein | sperm membrane | required for fertilization upon gamete contact |
| *spe-9* | *spe-9(hc88) I* | BA671 | **fertilization** | EGF repeat-containing protein | near sperm membrane | required for fertilization upon gamete contact |
| *spv-1* | *spv-1(ok1498) II* | RB1353 | **ovulation** | F-BAR/RhoGAP protein | membranes of neurons, spermatheca, hypodermis | required for normal spermathecal contractions |
| *vab-1* | *vab-1(dx31) II* | CZ337 | **attraction + ovulation** | ephrin receptor | neurons, oocytes, sheath cells | required for normal sperm migration toward oocytes, sheath cell contractions; receptor for MSP |
| *vab-1; fog-2* | *vab-1(dx31)/mIn1 [dpy-10(e128) mIs14] II; fog-2(q71) V* | DG1612 | **attraction + ovulation** | ephrin receptor | neurons, oocytes, sheath cells | required for normal sperm migration toward oocytes, sheath cell contractions; receptor for MSP |

## Supplementary Figures

**Supplementary Figure S1.**
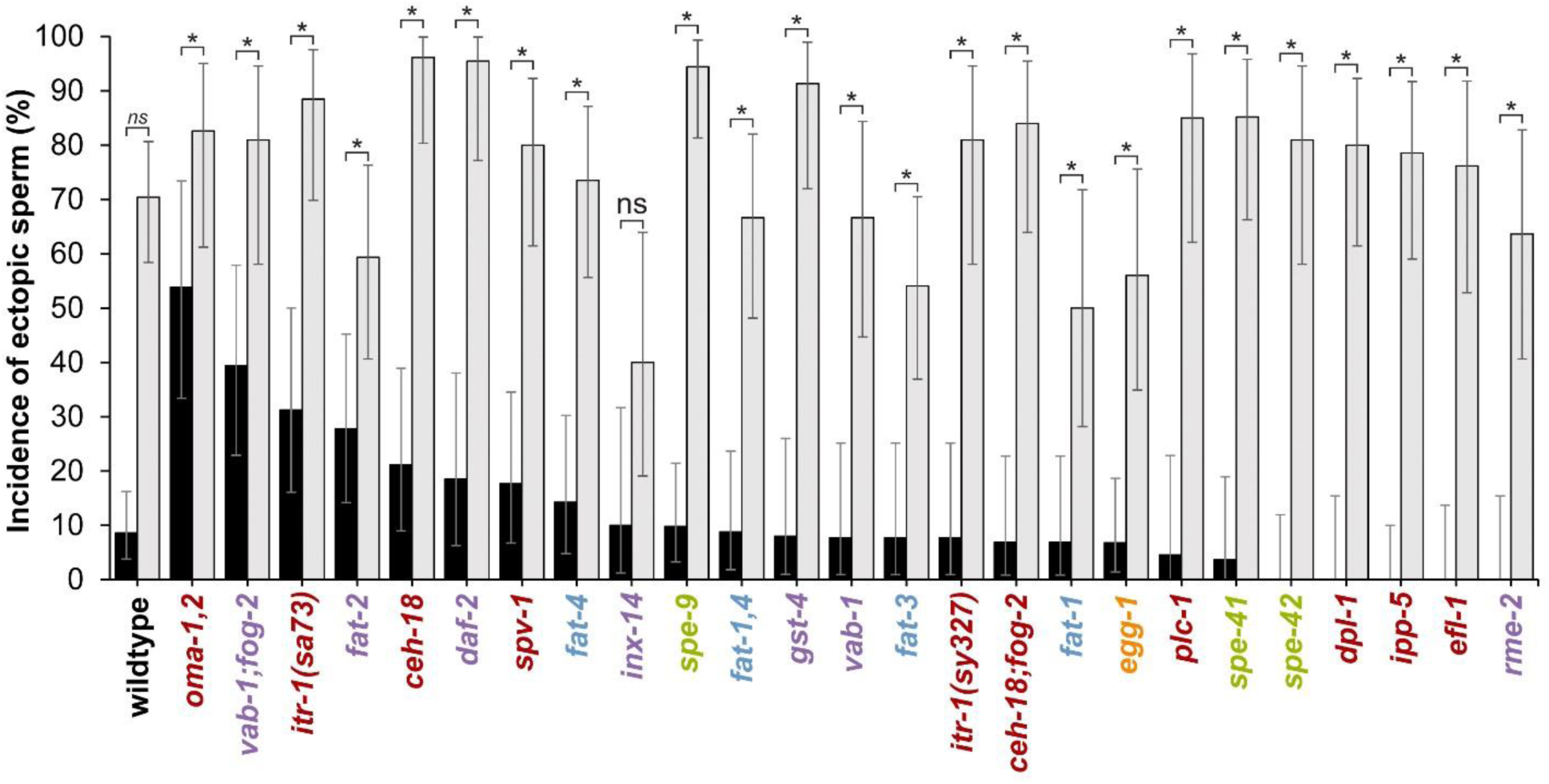
Mutant genotypes of *Caenorhabditis elegans* differ in the incidence of sperm invasion from heterospecific and conspecific male sperm. Incidence of ectopic sperm differs significantly for all strains when mated to conspecific *C. elegans* males (black bars) versus heterospecific *C. nigoni* males (gray bars) (except *inx*-*14*). Asterisks indicate statistical significance (P≤0.05) from χ^2^-tests on a given mutant strain (ns indicates non-significant difference P>0.05). Mutant strain names are colored to indicate functional phenotypic effects as in Figure 1 in the main text.

**Supplementary Figure S2.**
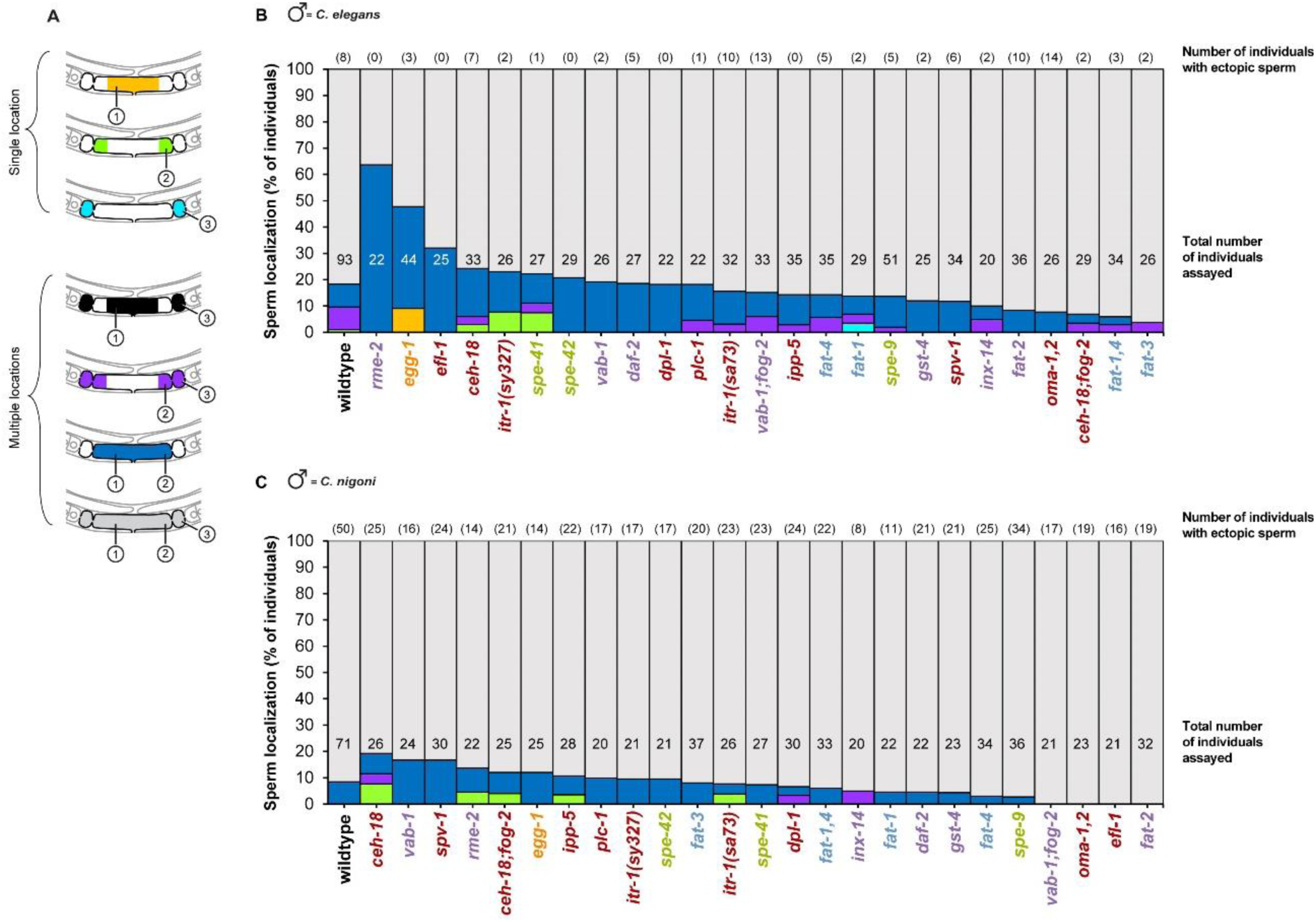
Sperm localization in non-ectopic regions. **(A)** Categories of sperm localization within non-ectopic regions of the reproductive tract, indicated for combinations of sperm present in proximal and distal uterus and spermatheca. **(B)** Relative incidence of conspecific *C. elegans* sperm in non-ectopic locations illustrates how sperm presence in all three regions is most common for nearly all mutant strains (gray) and differs significantly across strains (χ^2^=262.9, df=125, P<0.0001). **(C)** Heterospecific *C. nigoni* male sperm also were found most commonly in all three non-ectopic regions of the reproductive tract and did not differ significantly across strains (χ^2^=82.1, df=75, P=0.27). Total sample size of the number of mated hermaphrodites assayed indicated toward the bottom of each bar; the subset of individuals that also exhibited ectopic sperm indicated above each bar in parentheses. Mutant strain names are colored to indicate functional phenotypic effects as in Figure 1 in the main text.

**Supplementary Figure S3.**
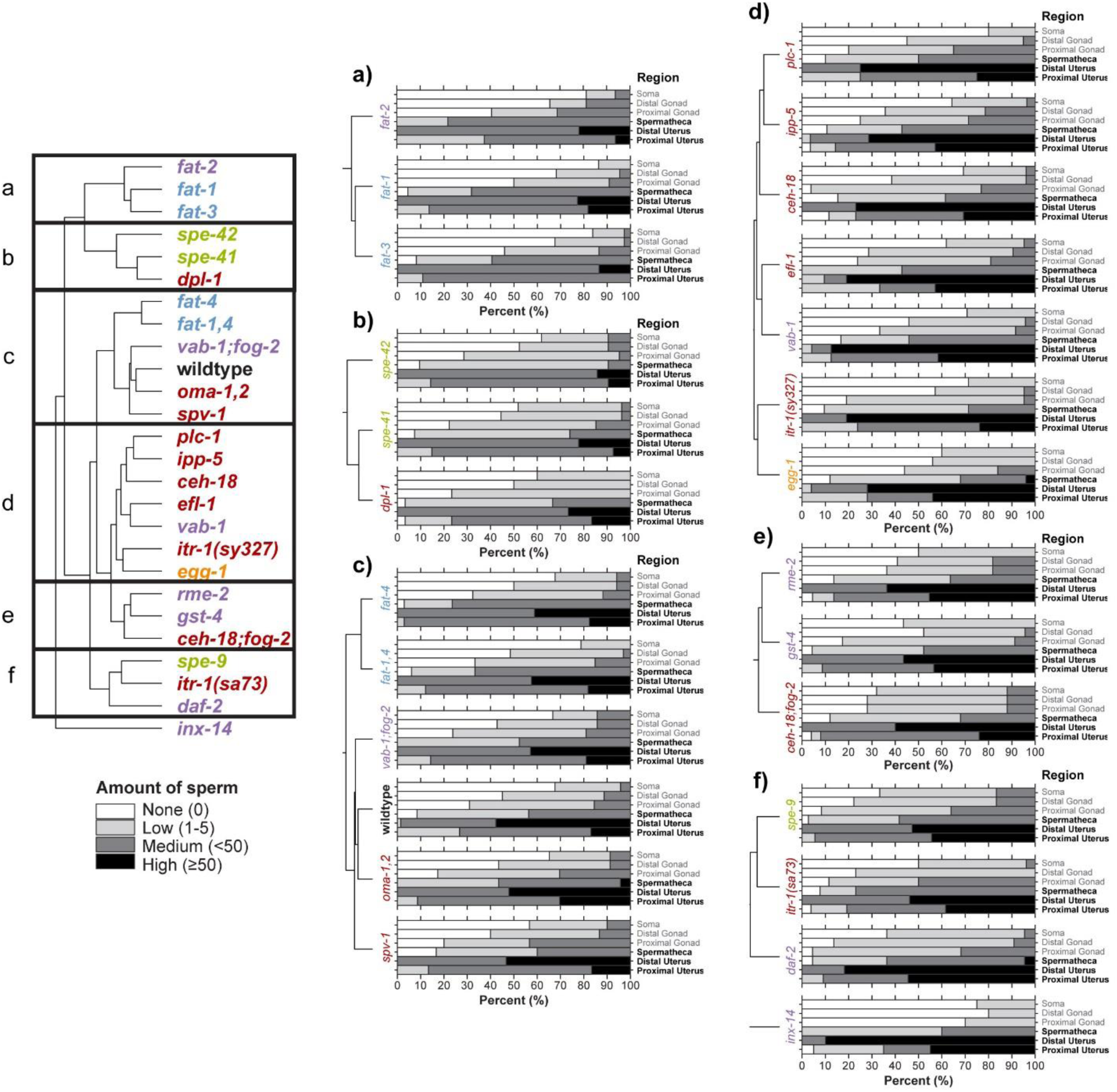
Distribution of heterospecific *C. nigoni* male sperm in the bodies of mated hermaphrodites. Stacked bars depict the cumulative relative abundance of heterospecific *C. nigoni* sperm observed in each region of the body as the proportion of mated hermaphrodites, shown separately for each mutant strain. Sperm abundance in each region for a given individual ranged from none (white), ‘low’ (1-5 sperm in the region, light gray), ‘medium’ (<50 sperm present, dark gray), to high (≥50 sperm present, black), assessed for the three non-ectopic regions and the three ectopic regions defined in Figure 1. Mutant strains are organized by the dendrogram of phenotypic similarity from Figure 4 as strain clusters (a-f). Ectopic regions are indicated in bold text. Mutant strain names are colored to indicate functional phenotypic effects as in Figure 1 in the main text.

**Supplementary Figure S4.**
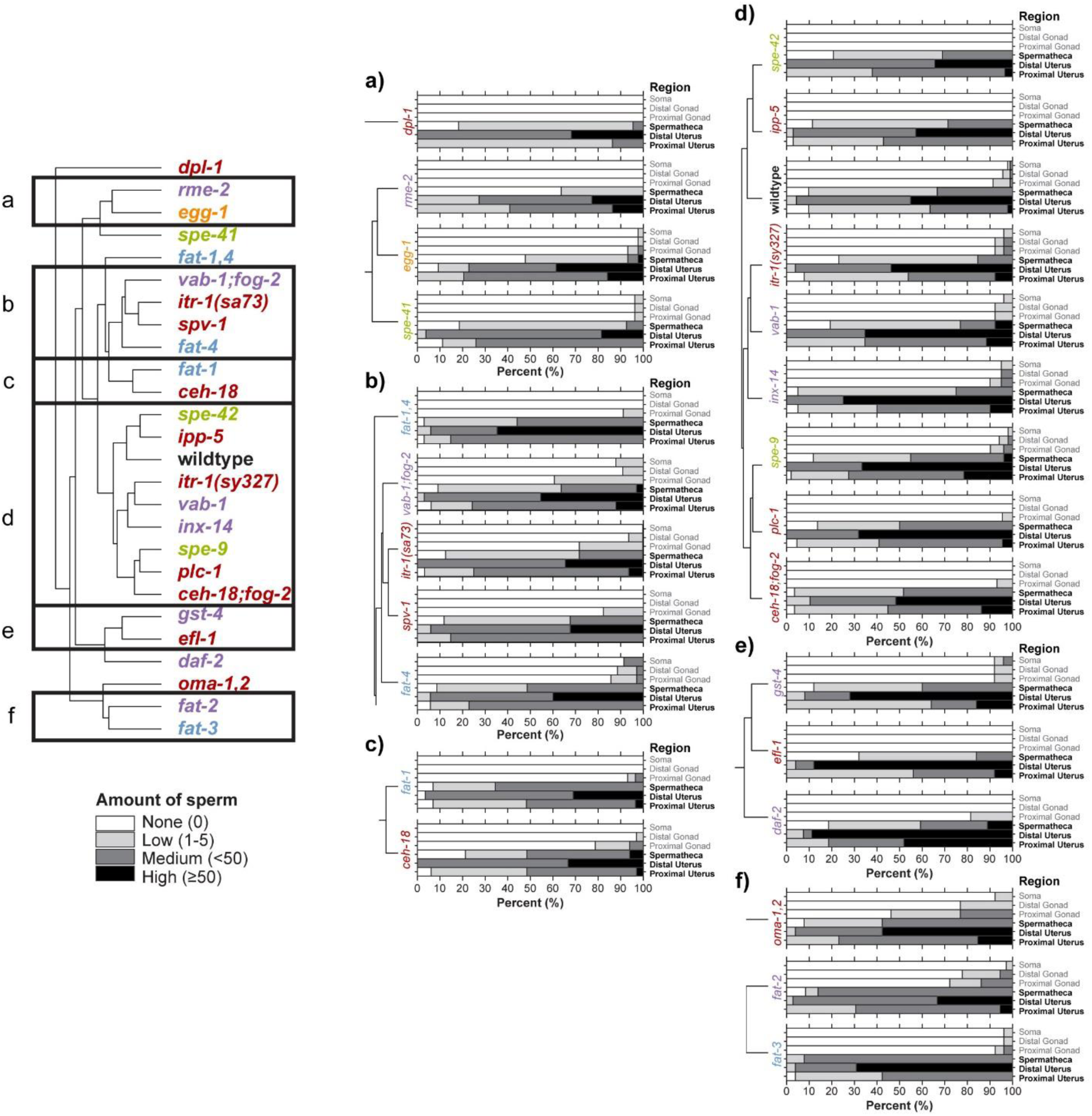
Distribution of conspecific *C. elegans* male sperm in the bodies of mated hermaphrodites. Stacked bars depict the cumulative relative abundance of conspecific *C. elegans* sperm observed in each region of the body as the proportion of mated hermaphrodites, shown separately for each mutant strain. Sperm abundance in each region for a given individual ranged from none (white), ‘low’ (1-5 sperm in the region, light gray), ‘medium’ (<50 sperm present, dark gray), to high (≥50 sperm present, black), assessed for the three non-ectopic regions and the three ectopic regions defined in Figure 1. Mutant strains are organized by the dendrogram of phenotypic similarity from Figure 4 as strain clusters (a-f). Ectopic regions are indicated in bold text. Mutant strain names are colored to indicate functional phenotypic effects as in Figure 1 in the main text.

